# An empirical evaluation of genotype imputation of ancient DNA

**DOI:** 10.1101/2021.12.22.473913

**Authors:** Kristiina Ausmees, Federico Sanchez-Quinto, Mattias Jakobsson, Carl Nettelblad

## Abstract

With capabilities of sequencing ancient DNA to high coverage often limited by sample quality or cost, imputation of missing genotypes presents a possibility to increase power of inference as well as cost-effectiveness for the analysis of ancient data. However, the high degree of uncertainty often associated with ancient DNA poses several methodological challenges, and performance of imputation methods in this context has not been fully explored. To gain further insights, we performed a systematic evaluation of imputation of ancient data using Beagle 4.0 and reference data from phase 3 of the 1000 Genomes project, investigating the effects of coverage, phased reference and study sample size. Making use of five ancient samples with high-coverage data available, we evaluated imputed data with respect to accuracy, reference bias and genetic affinities as captured by PCA. We obtained genotype concordance levels of over 99% for data with 1x coverage, and similar levels of accuracy and reference bias at levels as low as 0.75x. Our findings suggest that using imputed data can be a realistic option for various population genetic analyses even for data in coverage ranges below 1x. We also show that a large and varied phased reference set as well as the inclusion of low- to moderate-coverage ancient samples can increase imputation performance, particularly for rare alleles. In-depth analysis of imputed data with respect to genetic variants and allele frequencies gave further insight into the nature of errors arising during imputation, and can provide practical guidelines for post-processing and validation prior to downstream analysis.

## INTRODUCTION

The possibility to sequence ancient DNA (aDNA) has increased capabilities to study archaeological remains and provided new insights into various aspects of human evolutionary history. Notable findings such as the detection of genetic introgression between anatomically modern humans and other hominins, confirmation of the African origin of modern humans, and an increased understanding of the spread of agriculture into Europe have been achieved through population genetic analyses of ancient and contemporary genomes (Nielsen *et al.* 2017).

Due to the age and varying preservation conditions that ancient samples may have been exposed to, aDNA has unique properties that pose methodological and computational challenges not present when working with data from present-day humans. Contamination of DNA from microbes and other non-target sources can result in low proportions of endogenous DNA (Pääbo *et al.* 2004; Prüfer *et al.* 2010), leading to limitations in sample availability that can cause sequencing to high coverage depth to be impossible or prohibitively expensive. Contamination also leads to issues regarding data authenticity. In addition, the degradation of the DNA molecule that occurs over time leads to damage in the form of fragmentation and nucleotide misincorporations (Sánchez-Quinto *et al.* 2012; Brotherton *et al.* 2007), which can result in shorter fragments and higher levels of sequencing errors.

Although the identification of patterns of damage unique to aDNA has allowed for methods of data authentication and improved the process of reconstructing ancient genomes (Sawyer *et al.* 2012; Stiller *et al.* 2006; Briggs *et al.* 2007; Krause *et al.* 2010), these characteristics nonetheless cause biases in sequencing and mapping that can impact variant calling and other forms of downstream analysis (Prüfer *et al.* 2010; Ginolhac *et al.* 2011; Parks and Lambert 2015). Consequently, studies of aDNA samples are often limited to low- to moderate-coverage data with higher degrees of uncertainty than modern samples typically exhibit.

Genotype imputation is a powerful tool that can increase the information content in a sample by inferring unobserved genotypes. Imputation has been widely applied in various scenarios analyzing modern data, e.g. to increase power of inference in genome-wide association studies and to conform samples from different studies for merged analysis (Marchini and Howie 2010; Spencer *et al.* 2009; Zeggini *et al.* 2008). For aDNA, the possibility to increase information content of sparse and noisy data can potentially improve the quality of results as well as expand the range of analyses that are possible to perform.

Many common imputation methods for unrelated samples rely on sequential probabilistic models in which missing genotypes are inferred based on similarity to other individuals. The estimation is founded on an assumption of the presence of short stretches of shared haplotypes that have been passed down from a distant common ancestor. Given the individual’s genotypes, its haplotype phase is inferred, allowing missing variants to be predicted based on similarity to other samples. Most methods are able to leverage the information in the study sample as well as an additional panel of phased reference haplotypes when performing the phase estimation.

Many widely employed tools, such as MACH (Li *et al.* 2010), IMPUTE2 (Howie *et al.* 2009) and PHASE (Stephens *et al.* 2001; Stephens and Scheet 2005), are based on variants of the ‘the product of approximate conditionals’ (PAC) framework (Li and Stephens 2003). This model represents the sample sequence as an imperfect mosaic of the reference haplotypes, generally considering all possible transitions over the state space, with explicit modeling of mutation and recombination. A haploid version of this framework is used in the software GLIMPSE (Rubinacci *et al.* 2020). A slightly different method is implemented in the software Beagle (Browning and Browning 2007), which is based on a model of local haplotype clusters based on similarity of the reference haplotypes at nearby markers. This results in a smaller state space with not all possible transitions considered at every position, reducing the computational burden. The effects of mutation and recombination are not explicitly modeled, but the change of cluster membership along the sequence can be seen as implicitly representing these biological processes.

The accuracy of genotype imputation depends on several factors, mainly related to the quality of the sample data and the properties of the phased reference panel. A larger sample size, as well as increased marker density and genotype accuracy generally results in better performance (Browning and Browning 2011). For data with high levels of uncertainty, imputation based on a probabilistic framework using genotype likelihoods rather than called genotypes may be beneficial (Browning and Browning 2011; Browning and Yu 2009; Nielsen *et al.* 2011), an option that is supported by some software tools, including Beagle v4.0. In Hui *et al.* (2020), a two-step approach is introduced, in which genotype likelihood data is first used to obtain genotype probabilities based on a reference panel, and missing genotypes are subsequently imputed based on a subset of these which were able to be confidently called. They evaluate different pipelines on low coverage data below 1x and show that applying the two-step method using Beagle v4.1 or GLIMPSE for calling genotypes and Beagle v5 for imputation gave similar overall accuracy as using the single-step methods Beagle v4.0 or GLIMPSE, but that the two-step method gave more nuanced posterior genotype probabilities which allowed for a more informed post-imputation filtering procedure.

Studies comparing different phased reference panels have yielded varying results, with some finding that highest performance is gained by using population-specific panels (Mitt *et al.* 2017; Pistis *et al.* 2015) and others indicating the benefits of a large reference with a high level of diversity, particularly for admixed population with no clearly matching reference (Jostins *et al.* 2011; Huang *et al.* 2009; Howie *et al.* 2011). Using a phased genome reference panel from present-day individuals is currently the only option for ancient data, introducing a possible source of bias as it means that only variants that exist in the population today can be reproduced. Leveraging information in other ancient individuals by increasing study sample size may be a way to mitigate reference divergence, particularly as more sequenced ancient genomes become available, but benefits may be diminished in the context of sparse and uncertain data. The behavior of genotype likelihood-based imputation methods on data that exhibits the characteristic properties of aDNA discussed above has not been fully explored, particularly in combination with low coverage levels below 1x.

Genotype imputation has previously been performed on ancient human data in e.g. Gamba *et al.* (2014); Jones *et al.* (2015); Martiniano *et al.* (2017); Antonio *et al.* (2019); Cassidy *et al.* (2020). In these studies, imputation was performed using Beagle v4.0 and used to maximize the information content of ancient samples and allow for analyses such as Runs of Homozygosity (RoH) that require dense, diploid genotypes. Performance evaluation was mainly done by comparison of genotypes imputed from masked data to corresponding high-coverage calls, and showed satisfactory concordance to motivate the use of imputed data for downstream population-genetic analyses. The goal of this study is to complement and extend previous work by performing a systematic evaluation of a commonly used genotype likelihood-based imputation pipeline on ancient data. We investigate how the particular issues of sample quality and reference divergence associated with aDNA affect imputation, focusing on practical considerations regarding methodology and performance evaluation.

## MATERIALS AND METHODS

### Data description and preprocessing

#### Ancient samples

The ancient genome data used in this study consisted of five individuals for which high-coverage data between 19x to 57x was available (ans17, LBK, Loschbour, sf12 and ne1), as well as a set of 61 samples with low- to moderate-coverage ranging from 0.1x to 16x. See Supplementary Table 1 and Supplementary Table 2 for sample specifics and references to source publications.

The Genome Analysis Toolkit (GATK) v3.5.0 (McKenna *et al.* 2010) tool UnifiedGenotyper was used to generate genotype likelihoods from alignment data for each of the ancient samples individually. The allele callset used was that of the 1000 Genomes phase 3 panel (Auton *et al.* 2015), filtered to keep only autosomal, biallelic SNPs, resulting in a total of 77818182 markers. In order to avoid introducing a possible bias from nucleotide misincorporations due to post-mortem damage, the generated VCF files were filtered to exclude all sites where the most likely genotype could have been inferred from a deaminated allele. For C→T deaminations, this was done by removing sites where the SNP was a C→T transition and the most likely genotype contained a T allele. The corresponding treatment was performed for G→A deaminations. The software bcftools v1.6 (Li *et al.* 2009) was used for filtering.

As in previous studies considering aDNA, assessment of imputation performance was done by comparison of imputation results to corresponding high-quality (HQ) genotypes. For this, the five samples for which high-coverage data was available were used. The HQ genotypes considered as gold standard were called from the original high-coverage data, following the same pipeline as described in their respective publications (Supplementary Table 1). The called genotypes were filtered to keep only biallelic SNPs with a minimum depth of 15 and a QUAL score of at least 50. Heterozygote sites were further filtered for both alleles having a minimum allele depth of 25% of the total depth. Low-coverage data for each evaluation sample was generated by downsampling reads using Picard version 2.0.1 (Broad Institute version 2.0.1), after which estimation of genotype likelihoods and filtering were performed in an identical manner to the method described above for the low- to moderate-coverage samples. The sparse data was used as input for imputation, and the resulting genotypes were compared to HQ data.

#### Reference panels

The reference material used for imputation was the 1000 Genomes phase 3 v5a panel of phased haplotypes and the GRCh37 genetic maps provided along with the Beagle software. The reference panel was filtered to only include biallelic SNPs, resulting in 27904756 markers over chromosomes 1-22. In order to evaluate the effects of the reference on imputation, two panels were considered: the entire data set of 2504 samples, and a smaller one of 254 samples from European populations only. The two panels are denoted as “FULL” and “EUR”, respectively. The former was also used to estimate Minor Allele Frequency (MAF) of SNPs when analysis of imputed data over the allele frequency spectrum was performed.

### Imputation methodology

Imputation was performed using Beagle v4.0, with sample data split into segments of 50,000 markers with an overlap of 25,000, using genotype likelihoods as input. Version 4.0 was selected as the goal was to evaluate imputation based on probabilistic input in the context of extremely low coverages, and this is the latest version of the software that allows imputation to be performed based on genotype likelihoods. Further, we also wanted to assess the effects of including other ancient samples in the panel, allowing the imputation to be informed by other low-coverage ancient samples within the same probabilistic framework. The effects of including multiple ancient samples in the panel were evaluated by performing imputation jointly as well as separately. In the first case, all 61 low-to-moderate coverage samples as well as the 5 evaluation samples were included in the panel to imputed. In the second case, imputation was performed separately per study individual, meaning only the phased reference haplotypes were used in the genotype estimation.

### Performance evaluation metrics

#### Genotype concordance

The main metric employed for assessing imputation accuracy was genotype concordance/discordance, defined as the fraction of genotypes that were imputed correctly/incorrectly. This was measured separately for each of the 5 evaluation samples by comparison to the HQ genotypes derived from dense data. For each of the five evaluation samples, we extracted the imputed markers for which there were corresponding HQ genotypes available, and divided them into two disjoint sets denoted as “overlapping” and “non-overlapping”, based on whether or not the corresponding downsampled individual data had overlapping reads for the site or not. Supplementary Table 3 shows the sizes of these sets for each evaluation sample and coverage level.

#### Reference affinity

To assess whether imputed genotypes show a systematic bias towards the reference panel, the degree to which a sample showed an affinity towards the reference was compared between imputed and HQ genotypes. Here, reference affinity was measured as the fraction of markers that have the same genotype as the most frequently occurring one in the reference panel.

#### PCA

Principal Component Analysis (PCA) is a method of projecting data onto a basis that maximizes the variance of the data, possibly revealing previously unseen patterns or features. PCA can be used to reduce the dimensionality of data, e.g. for visualization purposes, and in the field of aDNA it is commonly used to show ancient samples in the context of modern variation. Here, it was used as a means of illustrating the difference between imputed and corresponding high-coverage genotypes.

PCA was performed on diploid genotypes, with a modern panel consisting of 429 European samples from the Human Origins data set of Patterson *et al.* (2012), filtered to remove variants with MAF under 1% or missing call rates exceeding 10%. In order to handle the fact that the ancient samples did not have observed genotypes for all sites used in the PCA, we used the method of Known Data Regression (KDR) (Arteaga and Ferrer 2002). A reference PCA based on the modern panel was initially defined. Estimation of scores for each ancient sample based on this model proceeded by considering the data of the reference samples corresponding to observed ancient genotypes, and fitting a linear regression model to their original PCA scores in the reference model. The software `SMARTPCA` from `EIGENSOFT 7.2.1` was used to define the reference PCA, and the Python library `scikit-learn` was used for solving linear least squares problems.

## RESULTS

### Effects of reference set and study sample size

Imputation performance was assessed for three combinations of reference set and study sample size, denoted as imputation configurations and shown in Table 1. In the first configuration, imputation was performed individually per evaluation sample, using the EUR phased reference set. For configurations 2 and 3, all ancient individuals were included in the imputation, using the EUR and FULL reference sets, respectively. All results in this section are for imputation performed on sample data with 1x coverage, and in order to perform a comprehensive evaluation of the effect of imputation configuration, no posterior filter was imposed on the imputed genotypes. Results are shown for each of the five evaluation samples, with results split into overlapping and non-overlapping marker sets.

**Table 1.** Imputation configurations.

| Configuration | Reference | Study Sample |
| --- | --- | --- |
| 1 | EUR | single |
| 2 | EUR | all ancient |
| 3 | FULL | all ancient |

Figure 1 shows genotype concordance for the three imputation configurations. Overall, concordance rates were similar between samples and reached 0.99 in all cases. The results indicate that the larger reference set as well as the inclusion of ancient samples in the imputation panel improved performance. For overlapping markers, concordance rates increased slightly from just under 0.997 to somewhat above. Concordance was lower among the non-overlapping markers in general, and it was also among these that the effects of varying imputation configurations were the most pronounced.

**Figure 1.**
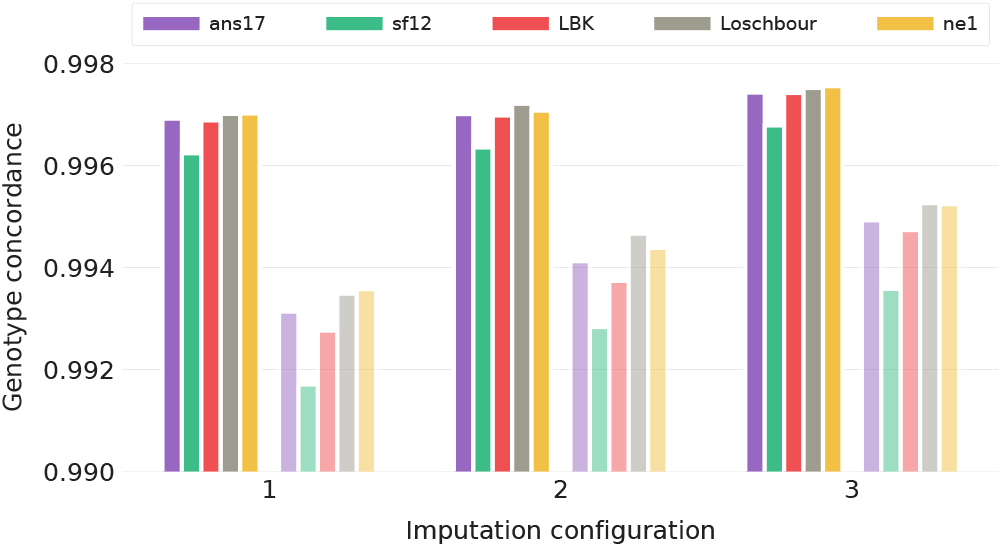
Concordance of imputed genotypes for the three configurations in Table 1. Imputation was performed on data in which the five evaluation individuals were downsampled to 1x coverage, and the evaluation was based on unfiltered results, with fully colored bars showing results for sites at which the downsampled data had overlapping reads, and shaded bars for non-overlapping markers, i.e. sites at which all overlapping reads were removed by the downsampling process.

In order to investigate imputation performance over the allele frequency range, the imputed markers were binned according to MAF, and the average genotype discordance assessed per bin.

Since there is a high risk of chance agreement between homozygous genotypes and the reference majority in the case of low MAF markers, only sites at which the HQ data was heterozygous were considered, thus measuring the ability of the imputation to recover heterozygotes. Results for imputation configurations 1-3 are shown in Figure 2, and again indicate that using a larger reference panel and study sample size increases performance. The effects were particularly visible among non-overlapping markers and in the low MAF ranges, where heterozygotes are the most difficult to recover.

**Figure 2.**
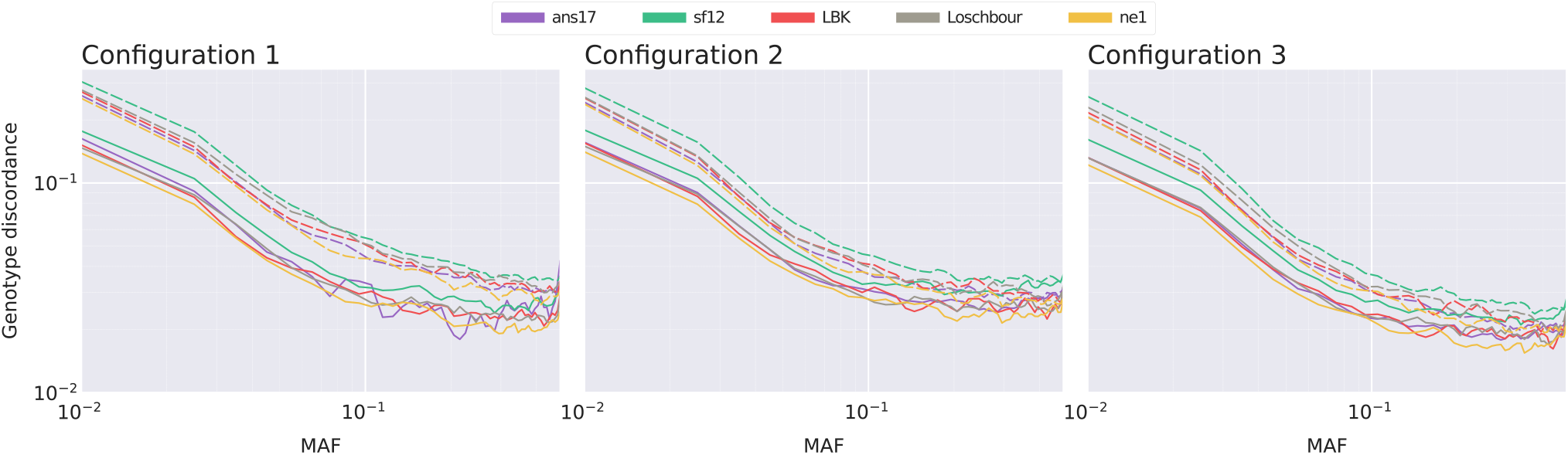
Log-log plots displaying discordance of heterozygote genotypes for the five evaluation samples. The subplots show results obtained using imputation configurations 1-3 described in Table 1, averaged over markers in 50 MAF bins. Results are shown for unfiltered imputed genotypes of data downsampled to 1x coverage, with solid lines indicating overlapping markers, and dashed lines markers at which the down-sampled data had no overlapping reads.

### Effects of coverage

Next, we assessed the effects of coverage on imputation. For every level *C* ∈ {0.10, 0.25, 0.50, 0.75, 1.00, 1.25, 1.50, 1.75, 2.00}, an imputation run was performed using the five evaluation samples downsampled to *C*x. Imputation configuration 2 was used for all runs, and as in the previous section, no posterior filter was imposed on the resulting genotypes.

Figure 3 shows the concordance between imputed and HQ genotypes for all markers (A) and for heterozygotes (B), separated into overlapping and non-overlapping marker sets for every evaluation sample and coverage level. A total concordance rate of 0.99 was reached around 0.25x for most samples, with a planing out visible at 1x where levels around 0.9975 and 0.995 were obtained for overlapping and non-overlapping sites, respectively. For heterozygote sites, concordance levels were below 0.975 throughout, reaching 0.95 at 1x and as low as 0.825 for the lowest coverage level of 0.1x.

**Figure 3.**
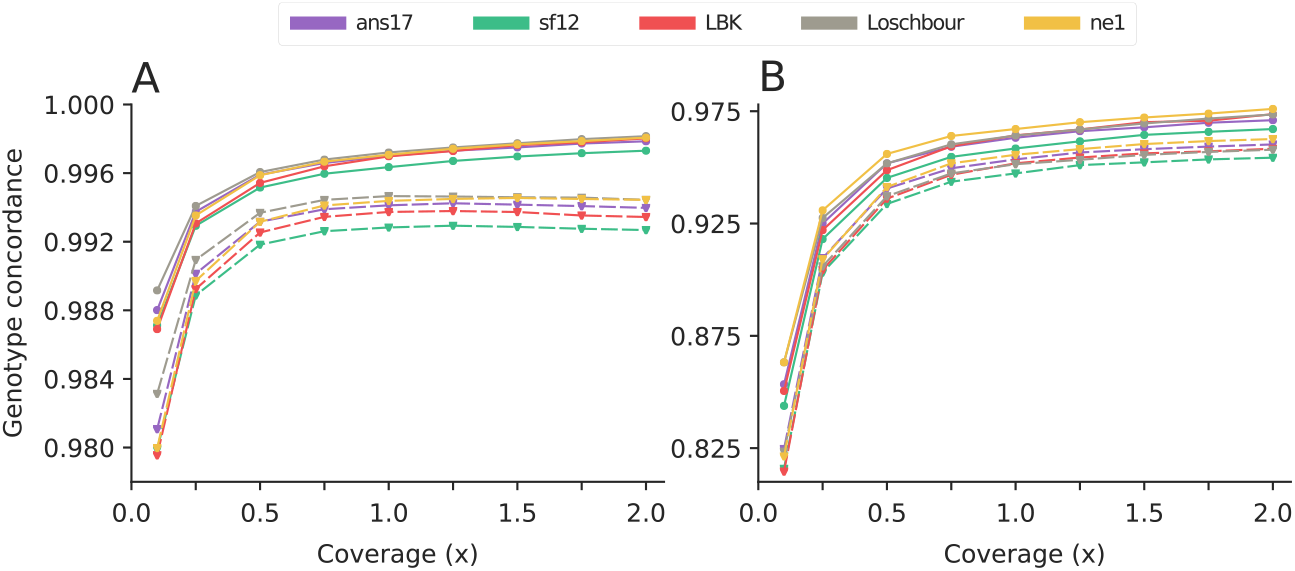
Genotype concordance of imputed data for different levels of coverage of the input data, for A: all markers and B: markers at which the HQ genotype was heterozygote. Results are shown for the five evaluation samples, with solid and dashed lines indicating SNPs with and without overlapping reads in the downsampled data, respectively. Imputation was performed using configuration 2 (Table 1), with no posterior filter imposed on the resulting genotypes.

In order to assess the level of systematic bias towards the variant that is most common in the reference, we compared the level of reference affinity of the imputed data to that of the corresponding HQ genotypes, considering these as a baseline for the similarity between the true genotypes and the reference. Figure 4 shows the difference in measured affinities between the HQ and imputed data for the 9 coverage levels, over the allele frequency spectrum, averaged over the 5 evaluation samples.

**Figure 4.**
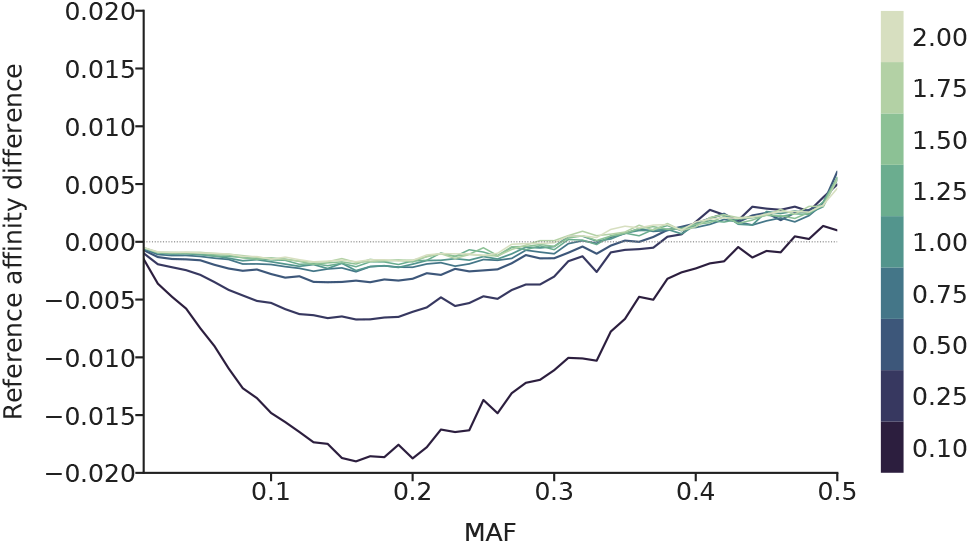
Comparison of levels of reference bias for different coverage levels. Input data for imputation in which the evaluation samples were downsampled to coverages ranging between 0.1-2.0x was generated (displayed in different colors), and the difference in reference affinity between HQ and resulting imputed genotypes estimated. Imputation was performed using configuration 2 (Table 1), with no posterior filter imposed on the resulting genotypes. Results are shown for all markers, aggregated into 50 MAF bins, and averaged over the evaluation samples.

The negative values in the lower MAF ranges indicate that the imputed data shows a larger affinity towards the reference. While the extremely low coverages showed significantly higher levels of bias, the rates decreased and showed little variation for coverages 0.75x and higher. The results indicate a reduction in bias towards the reference with increasing MAF, showing little differences around MAF 0.3 for most coverage levels. A possible explanation for the positive difference at higher MAF values is that at these markers, imputation errors do not tend towards the reference majority as strongly as in the lower MAF ranges. Overall concordance is also lower at high MAF due to higher frequency of heterozygotes.

### In-depth performance analysis

In this section we present further evaluation of imputed genotypes to assess properties relevant to downstream analysis. We considered results of imputation configuration 3 on data with 1x coverage, and imposed a filter of minimum posterior genotype probability of 0.99 on the imputed data. First, performance for different genotypes was evaluated. Figure 5 shows concordance of sites split according genotype in the high-coverage data. Although performance remained poorer for heterozygote sites, the filtered data showed improved levels of over 0.99 throughout. The filtered results also showed little difference between overlapping and non-overlapping marker sets at homozygote sites, while larger differences remained for heterozygotes. Inspection of performance for heterozygote sites across the allele frequency spectrum showed that discordance levels below 0.01 were reached around MAF 0.1, with over 85% of sites retained post-filter (Figure 6).

**Figure 5.**
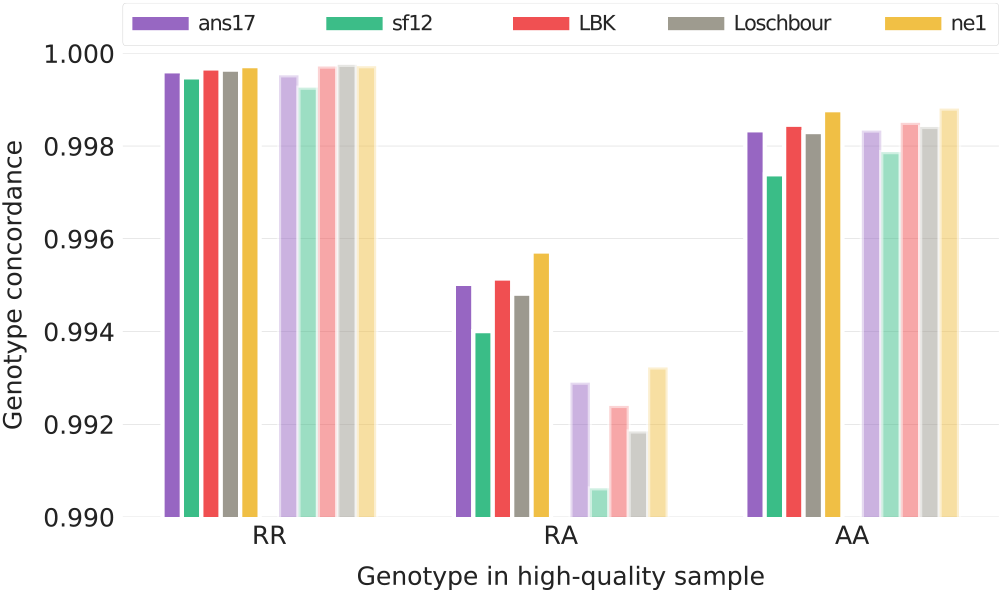
Concordance of imputed genotypes, split according to genotype in the HQ data. Imputation was performed on data in which the five evaluation individuals were downsampled to 1x coverage, using imputation configuration 3 (Table 1), and the resulting data was filtered for minimum genotype probability of 0.99. Fully colored bars indicate markers that had overlapping reads in the downsampled data, and shaded bars indicate sites that did not.

**Figure 6.**
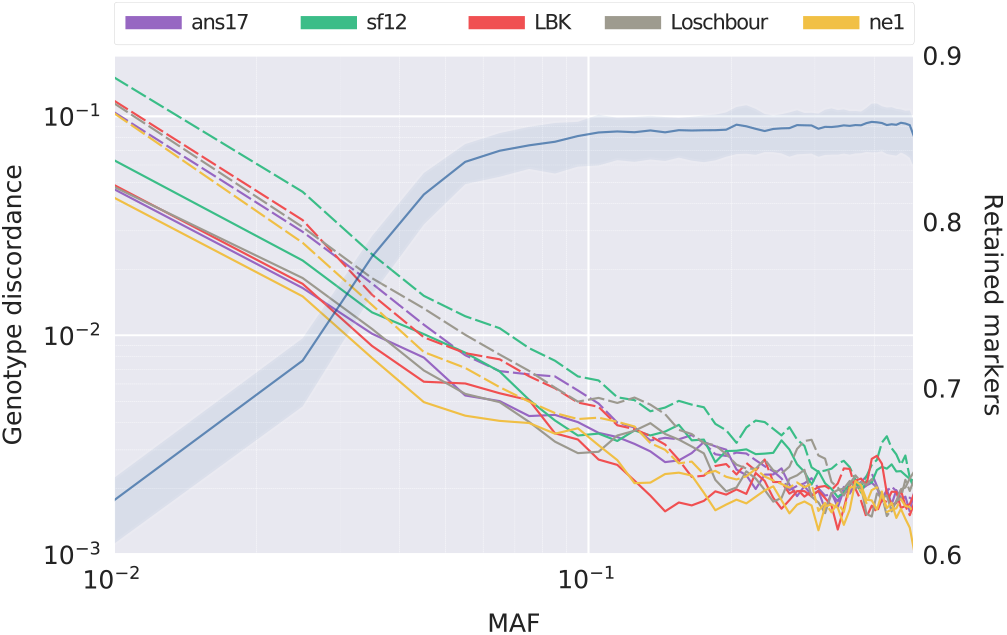
Log-log plot of genotype discordance at heterozygote sites, averaged over 50 bins in the allele frequency spectrum and smoothed using a moving average using 3 points. Input data for imputation was downsampled to 1x and configuration 3 (Table 1) was used, after which a posterior filter of minimum genotype probability of 0.99 was applied. The fraction of markers retained after the filter, averaged over the five evaluation samples, is shown in blue, with shaded regions indicating minimum and maximum. Solid and dashed lines indicate SNPs with and without overlapping reads in the downsampled data, respectively.

Finally, PCA was used to visualize and compare genetic affinities of imputed and high-coverage data. Figure 7 shows that, within the variation represented by the first two principal components, scores of imputed genotypes map closely to those of the HQ data, particularly for the three samples ans17, LBK and ne1. As illustrated in e.g. Günther and Jakobsson (2019), considering subsets of SNPs introduces noise in the PCA projection, resulting in less accurate projections with higher degree of uncertainty compared to using all available data. In Figure 7, this is visualized by the distance between low-coverage and HQ data points for each individual. For each of the five samples, relative distances to the HQ data are smaller for the imputed genotypes than the low-coverage data, indicating that information lost by the downsampling process has been retained by imputation.

**Figure 7.**
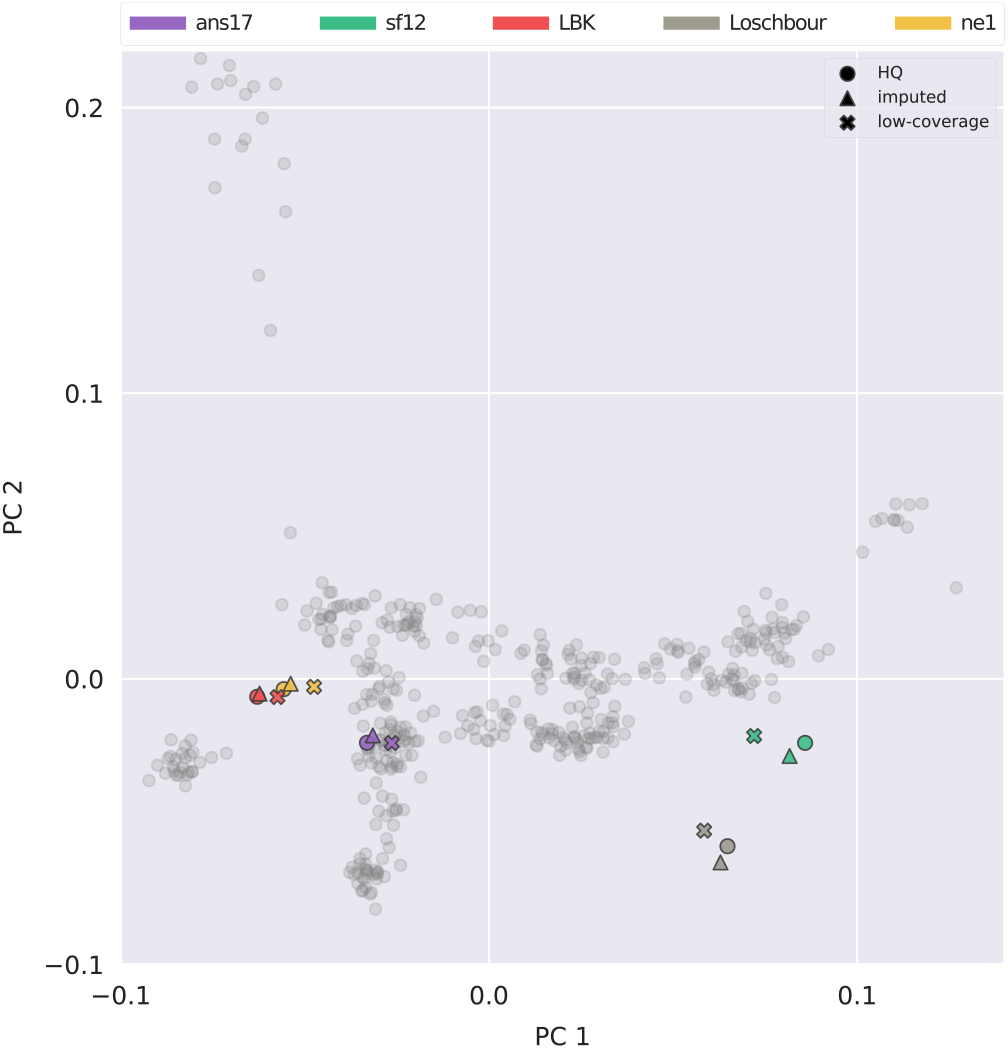
PCA comparing HQ, imputed and low-coverage data for the five evaluation individuals. A reference PCA was defined based on genotypes of modern European samples from the Human Origins (Patterson *et al.* 2012) data set, after which the KDR method was used to estimate scores of ancient samples. Modern individuals are indicated by gray dots and ancient samples coloured according to the legend. The low-coverage data points correspond to the genotypes that has been downsampled to 1x, filtered, and used as input to imputation, which was performed using configuration 3 (Table 1), with a posterior filter of minimum genotype probability of 0.99 applied.

## DISCUSSION

This study provides a systematic investigation of the application of imputation to human aDNA. We have corroborated results from similar experiments showing that overall genotype concordance levels of over 0.99 can be reached for data with 1x coverage, and provided an in-depth analysis of the qualities of imputed genotypes. Investigation of performance at coverage levels tending to the ultra-low indicated that the quality of imputed genotypes began to plateau around 0.75-1x, and that common variants were more robust to different reference sets as well as less prone to reference bias overall. The results suggest a MAF threshold of at least 0.1 for reliable heterozygote calls, and that in cases where rare alleles are of interest, an increased diversity and size of the phased reference as well as the imputation panel are particularly beneficial. The fact that information from ancient samples can be leveraged to improve imputation, along with the genotype concordance levels shown, may in some cases motivate the sequencing of more ancient samples to lower coverage as a cost-effective alternative to that of fewer samples to higher coverage.

The presented work provides a framework of practical considerations for performing imputation, as well as a basis for further investigations. A systematic evaluation of different imputation methods and adaptation of the statistical models to the context of sparse and uncertain data may increase performance for ancient samples. Further insights may also be gained by assessment of imputed data by means of different population genetic analyses, considering e.g. haplotype-based methods such as RoH as well as those based on allele frequencies.

For the five evaluation samples considered, in-depth performance analysis showed lower performance for the two hunter-gatherer genomes sf12 and Loschbour, both in terms of genotype concordance and similarity in PCA-space to HQ data. A possible explanation is that the reference panel of present-day individuals used for imputation does not contain individuals with a genetic composition that is similar to these samples. As discussed in e.g. Skoglund *et al.* (2014) and Günther and Jakobsson (2016), hunter-gatherer samples have shown a particular genetic profile that is not represented in the genetic variation of modern-day people. The samples LBK, ne1 and ans17, in contrast, are from early farming cultures, for which there are present-day European individuals who share a similar genetic make-up. Further investigations of the effects of sample ancestry and reference divergence will be required to customize imputation pipelines to samples of varying genetic composition. As more sequenced ancient samples become available, both imputation accuracy as well as the ability to assess various aspects of performance will be improved.

We have focused on Beagle v4.0 as it is a probabilistic imputation framework that has been frequently used in the aDNA community. Other imputation pipelines have shown to give qualitatively similar results for imputing genotypes from ancient sequence data, and also indicated some performance trade-offs such as imputation accuracy at different parts of the allele frequency spectrum (Hui *et al.* 2020). Another relevant point is that more recent software for phasing and imputation such as GLIMPSE and Beagle v5 have increased focus on scalability to larger reference panels. Sample size, availability of computational resources and the type of downstream analysis intended are thus factors to consider in the selection of methodology for imputation of aDNA.

## DATA AVAILABILITY

The authors state that all data necessary for confirming the conclusions presented in the article are represented fully within the article and in its online supplementary material.

## ACKNOWLEDGMENTS

The authors acknowledge the use of computational resources provided by Swedish National Infrastructure for Computing (SNIC) at Uppsala Multidisciplinary Center for Advanced Computational Science (UPPMAX) under project SNIC 2017/7-162. CN also acknowledges funding by Formas (grant number 2020-00712), and MJ acknowledges the Knut and Alice Wallenberg foundation.

## LITERATURE CITED

Antonio, M. L., Z. Gao, H. M. Moots, M. Lucci, F. Candilio, et al., 2019 Ancient rome: A genetic crossroads of europe and the mediterranean. Science 366: 708–714.

Arteaga, F. and A. Ferrer, 2002 Dealing with missing data in mspc: several methods, different interpretations, some examples. Journal of Chemometrics 16: 408–418.

Auton, A., G. R. Abecasis, D. M. Altshuler, R. M. Durbin, D. R. Bentley, et al., 2015 A global reference for human genetic variation. Nature 526: 68–74.

Briggs, A. W., U. Stenzel, P. L. F. Johnson, R. E. Green, J. Kelso, et al., 2007 Patterns of damage in genomic dna sequences from a neandertal. Proc Natl Acad Sci U S A 104: 14616–14621.

Broad Institute, version 2.0.1 Picard tools. http://broadinstitute.github.io/picard/.

Brotherton, P., P. Endicott, J. J. Sanchez, M. Beaumont, R. Barnett, et al., 2007 Novel high-resolution characterization of ancient dna reveals c > u-type base modification events as the sole cause of post mortem miscoding lesions. Nucleic Acids Res 35: 5717–5728.

Browning, B. L. and Z. Yu, 2009 Simultaneous genotype calling and haplotype phasing improves genotype accuracy and reduces false-positive associations for genome-wide association studies. Am J Hum Genet 85: 847–861.

Browning, S. R. and B. L. Browning, 2007 Rapid and accurate haplotype phasing and missing-data inference for whole-genome association studies by use of localized haplotype clustering. The American Journal of Human Genetics 81: 1084 – 1097.

Browning, S. R. and B. L. Browning, 2011 Haplotype phasing: existing methods and new developments. Nature Reviews Genetics 12: 703–714.

Cassidy, L. M., R. Ó. Maoldúin, T. Kador, A. Lynch, C. Jones, et al., 2020 A dynastic elite in monumental neolithic society. Nature 582: 384–388.

Gamba, C., E. R. Jones, M. D. Teasdale, R. L. McLaughlin, G. Gonzalez-Fortes, et al., 2014 Genome flux and stasis in a five millennium transect of european prehistory. Nature Communications 5: 5257.

Ginolhac, A., M. Rasmussen, M. T. P. Gilbert, E. Willerslev, and L. Orlando, 2011 mapdamage: testing for damage patterns in ancient dna sequences. Bioinformatics 27: 2153–2155.

Günther, T. and M. Jakobsson, 2016 Genes mirror migrations and cultures in prehistoric europe—a population genomic perspective. Current Opinion in Genetics Development 41: 115–123, Genetics of human origin.

Günther, T. and M. Jakobsson, 2019 Population genomic analyses of dna from ancient remains. In Handbook of statistical genomics, edited by D. J. Balding, I. Moltke, and J. Marioni, chapter 10, pp. 295–324, Wiley, Hoboken, NJ.

Howie, B., J. Marchini, and M. Stephens, 2011 Genotype imputation with thousands of genomes. G3 (Bethesda) 1: 457–470.

Howie, B. N., P. Donnelly, and J. Marchini, 2009 A flexible and accurate genotype imputation method for the next generation of genome-wide association studies. PLoS Genet 5: 1–15.

Huang, L., Y. Li, A. B. Singleton, J. A. Hardy, G. Abecasis, et al., 2009 Genotype-imputation accuracy across worldwide human populations. Am J Hum Genet 84: 235–250.

Hui, R., E. D’Atanasio, L. M. Cassidy, C. L. Scheib, and T. Kivisild, 2020 Evaluating genotype imputation pipeline for ultra-low coverage ancient genomes. Scientific Reports 10: 18542.

Jones, E. R., G. Gonzalez-Fortes, S. Connell, V. Siska, A. Eriksson, et al., 2015 Upper palaeolithic genomes reveal deep roots of modern eurasians. Nature Communications 6: 8912.

Jostins, L., K. I. Morley, and J. C. Barrett, 2011 Imputation of low-frequency variants using the hapmap3 benefits from large, diverse reference sets. Eur J Hum Genet 19: 662–666.

Krause, J., A. W. Briggs, M. Kircher, T. Maricic, N. Zwyns, et al., 2010 A complete mtdna genome of an early modern human from kostenki, russia. Current Biology 20: 231–236.

Li, H., B. Handsaker, A. Wysoker, T. Fennell, J. Ruan, et al., 2009 The sequence alignment/map format and samtools. Bioinformatics 25: 2078–2079.

Li, N. and M. Stephens, 2003 Modeling linkage disequilibrium and identifying recombination hotspots using single-nucleotide polymorphism data. Genetics 165: 2213–2233.

Li, Y., C. J. Willer, J. Ding, P. Scheet, and G. R. Abecasis, 2010 Mach: Using sequence and genotype data to estimate haplotypes and unobserved genotypes. Genet Epidemiol 34: 816–834.

Marchini, J. and B. Howie, 2010 Genotype imputation for genome-wide association studies. Nature Reviews Genetics 11: 499–511.

Martiniano, R., L. M. Cassidy, R. Ó’Maoldúin, R. McLaughlin, N. M. Silva, et al., 2017 The population genomics of archaeological transition in west iberia: Investigation of ancient substructure using imputation and haplotype-based methods. PLOS Genetics 13: 1–24.

McKenna, A., M. Hanna, E. Banks, A. Sivachenko, K. Cibulskis, et al., 2010 The genome analysis toolkit: a mapreduce framework for analyzing next-generation dna sequencing data. Genome Res 20: 1297–1303.

Mitt, M., M. Kals, K. Pärn, S. B. Gabriel, E. S. Lander, et al., 2017 Improved imputation accuracy of rare and low-frequency variants using population-specific high-coverage wgs-based imputation reference panel. Eur J Hum Genet 25: 869–876.

Nielsen, R., J. M. Akey, M. Jakobsson, J. K. Pritchard, S. Tishkoff, et al., 2017 Tracing the peopling of the world through genomics. Nature 541: 302–310.

Nielsen, R., J. S. Paul, A. Albrechtsen, and Y. S. Song, 2011 Geno-type and snp calling from next-generation sequencing data. Nature Reviews Genetics 12: 443–451.

Parks, M. and D. Lambert, 2015 Impacts of low coverage depths and post-mortem dna damage on variant calling: a simulation study. BMC Genomics 16: 19.

Patterson, N., P. Moorjani, Y. Luo, S. Mallick, N. Rohland, et al., 2012 Ancient admixture in human history. Genetics 192: 1065–1093.

Pistis, G., E. Porcu, S. I. Vrieze, C. Sidore, M. Steri, et al., 2015 Rare variant genotype imputation with thousands of study-specific whole-genome sequences: implications for cost-effective study designs. Eur J Hum Genet 23: 975–983.

Prüfer, K., U. Stenzel, M. Hofreiter, S. Pääbo, J. Kelso, et al., 2010 Computational challenges in the analysis of ancient dna. Genome Biol 11: R47–R47.

Pääbo, S., H. Poinar, D. Serre, V. Jaenicke-Després, J. Hebler, et al., 2004 Genetic analyses from ancient dna. Annual Review of Genetics 38: 645–679.

Rubinacci, S., D. Ribeiro, R. Hofmeister, and O. Delaneau, 2020 Efficient phasing and imputation of low-coverage sequencing data using large reference panels. bioRxiv.

Sawyer, S., J. Krause, K. Guschanski, V. Savolainen, and S. Pääbo, 2012 Temporal patterns of nucleotide misincorporations and dna fragmentation in ancient dna. PLoS One 7: 1–7.

Skoglund, P., H. Malmström, A. Omrak, M. Raghavan, C. Valdiosera, et al., 2014 Genomic diversity and admixture differs for stone-age scandinavian foragers and farmers. Science 344: 747–750.

Spencer, C. C. A., Z. Su, P. Donnelly, and J. Marchini, 2009 Designing genome-wide association studies: Sample size, power, imputation, and the choice of genotyping chip. PLoS Genet 5: 1–13.

Stephens, M. and P. Scheet, 2005 Accounting for decay of linkage disequilibrium in haplotype inference and missing-data imputation. Am J Hum Genet 76: 449–462.

Stephens, M., N. J. Smith, and P. Donnelly, 2001 A new statistical method for haplotype reconstruction from population data. Am J Hum Genet 68: 978–989.

Stiller, M., R. E. Green, M. Ronan, J. F. Simons, L. Du, et al., 2006 Patterns of nucleotide misincorporations during enzymatic amplification and direct large-scale sequencing of ancient dna. Proc Natl Acad Sci U S A 103: 13578–13584.

Sánchez-Quinto, F., H. Schroeder, O. Ramirez, M. Ávila Arcos, M. Pybus, et al., 2012 Genomic affinities of two 7,000-year-old iberian hunter-gatherers. Current Biology 22: 1494 – 1499.

Zeggini, E., L. J. Scott, R. Saxena, B. F. Voight, J. L. Marchini, et al., 2008 Meta-analysis of genome-wide association data and large-scale replication identifies additional susceptibility loci for type 2 diabetes. Nature Genetics 40: 638–645.

