## Supplementary material for "An empirical evaluation of genotype imputation of ancient DNA"

#### Data

| **High-coverage samples** | | | |
|---|---|---|---|
| | **Sample** | **Coverage (approx)** | **Reference** |
| 1 | Loschbour | 22 | Lazaridis et al. 2014 |
| 2 | LBK | 19 | Lazaridis et al. 2014 |
| 3 | ans17 | 27 | Fraser and Sánchez-Quinto (in preparation) |
| 4 | ne1 | 22 | Gamba et al. 2014 |
| 5 | sf12 | 57 | Günther et al. 2018 |

Table 1: The high-coverage samples used for performance evaluation.

| **Low- to moderate-coverage samples** | | | |
|---|---|---|---|
| | **Sample** | **Coverage (approx)** | **Reference** |
| 1 | BA64 | 9.68 | Cassidy et al. 2016 |
| 2 | Gok2 | 1.19 | Skoglund et al. 2014 |
| 3 | LC41 | 0.95 | Martiniano et al. 2017 |
| 4 | LC42 | 2.60 | Martiniano et al. 2017 |
| 5 | LC44 | 1.77 | Martiniano et al. 2017 |
| 6 | CA122A | 1.52 | Martiniano et al. 2017 |
| 7 | CM9B | 2.56 | Martiniano et al. 2017 |
| 8 | DOLDA96B | 1.69 | Martiniano et al. 2017 |
| 9 | CB13 | 0.85 | Olalde et al. 2015 |
| 10 | Iceman | 4.44 | Keller et al. 2012 |

| | | | |
|---|---|---|---|
| 11 | ne5 | 0.71 | Gamba et al. 2014 |
| 12 | ne6 | 0.86 | Gamba et al. 2014 |
| 13 | ne7 | 0.83 | Gamba et al. 2014 |
| 14 | CO1 | 0.79 | Gamba et al. 2014 |
| 15 | Bar8 | 6.28 | Hofmanová et al. 2016 |
| 16 | Bar31 | 3.37 | Hofmanová et al. 2016 |
| 17 | ATP2 | 8.71 | Günther et al. 2015 |
| 18 | ATP16 | 12.98 | Günther et al. 2015 |
| 19 | ATP12 | 2.43 | Günther et al. 2015 |
| 20 | mur | 3.33 | Valdiosera et al. 2018 |
| 21 | ans8 | 1.94 | Sánchez-Quinto et al. 2019 |
| 22 | ans14 | 2.58 | Sánchez-Quinto et al. 2019 |
| 23 | ajv70 | 1.34 | Günther et al. 2018 |
| 24 | ajv58 | 2.68 | Günther et al. 2018 |
| 25 | prs2 | 1.16 | Sánchez-Quinto et al. 2019 |
| 26 | prs9 | 1.89 | Sánchez-Quinto et al. 2019 |
| 27 | prs13 | 1.56 | Sánchez-Quinto et al. 2019 |
| 28 | prs16 | 1.78 | Sánchez-Quinto et al. 2019 |
| 29 | bal4 | 1.54 | Sánchez-Quinto et al. 2019 |
| 30 | kol6 | 1.48 | Sánchez-Quinto et al. 2019 |
| 31 | CO1CP | 0.17 | Mathieson et al. 2015 |
| 32 | ne1CP | 0.08 | Mathieson et al. 2015 |
| 33 | ne6CP | 0.21 | Mathieson et al. 2015 |
| 34 | ne7CP | 0.20 | Mathieson et al. 2015 |
| 35 | Motala12CP | 0.30 | Mathieson et al. 2015 |
| 36 | BranaCP | NA | Mathieson et al. 2015 |
| 37 | KO1CP | 0.20 | Mathieson et al. 2015 |
| 38 | Kotias | 15.40 | Jones et al. 2015 |
| 39 | Satsurblia | 2.16 | Jones et al. 2015 |
| 40 | Motala12SG | 1.94 | Lazaridis et al. 2014 |
| 41 | Bichon | 13.52 | Jones et al. 2015 |
| 42 | KO1 | 0.94 | Gamba et al. 2014 |
| 43 | LaBrana | 2.78 | Olalde et al. 2014 |
| 44 | Paliambela | 1.22 | Hofmanová et al. 2016 |
| 45 | Kleitos | 1.92 | Hofmanová et al. 2016 |
| 46 | Revenia | 1.02 | Hofmanová et al. 2016 |
| 47 | LatH1 | 0.86 | Jones et al. 2017 |
| 48 | LatH2 | 2.70 | Jones et al. 2017 |
| 49 | LatH3 | 0.62 | Jones et al. 2017 |
| 50 | LatMN1 | 0.12 | Jones et al. 2017 |
| 51 | Canes1 | 0.87 | González-Fortes et al. 2017 |
| 52 | SC1 | 0.98 | González-Fortes et al. 2017 |
| 53 | SC2 | 2.70 | González-Fortes et al. 2017 |

| | | | |
|---|---|---|---|
| 54 | OC1 | 1.51 | González-Fortes et al. 2017 |
| 55 | H26 | 4.00 | Günther et al. 2018 |
| 56 | SF9 | 1.15 | Günther et al. 2018 |
| 57 | sbj | 0.43 | Günther et al. 2018 |
| 58 | H22 | 0.71 | Günther et al. 2018 |
| 59 | steigen | 1.24 | Günther et al. 2018 |
| 60 | Kunila2 | 0.31 | Mittnik et al. 2018 |
| 61 | Gyvakarai1 | 2.00 | Mittnik et al. 2018 |

Table 2: The low- to moderate-coverage samples included in the imputation panel in configurations 2 and 3.

| Number of markers used in performance evaluation | | | | | | |
|---|---|---|---|---|---|---|
| | | **ans17** | **sf12** | **LBK** | **Loschbour** | **ne1** |
| total | | 26317153 | 27072696 | 15907150 | 17655877 | 19148120 |
| 0.1x | overlap | 2064457 | 2054700 | 1437112 | 1516225 | 1516936 |
| | no overlap | 24252696 | 25017996 | 14470038 | 16139652 | 17631184 |
| 0.25x | overlap | 4728775 | 4697958 | 3243931 | 3440090 | 3454067 |
| | no overlap | 21588378 | 22374738 | 12663219 | 14215787 | 15694053 |
| 0.5x | overlap | 8223690 | 8159398 | 5535230 | 5888457 | 5974794 |
| | no overlap | 18093463 | 18913298 | 10371920 | 11767420 | 13173326 |
| 0.75x | overlap | 10816510 | 10737175 | 7147381 | 7629381 | 7801581 |
| | no overlap | 15500643 | 16335521 | 8759769 | 10026496 | 11346539 |
| 1.0x | overlap | 12738211 | 12661508 | 8282383 | 8871611 | 9136265 |
| | no overlap | 13578942 | 14411188 | 7624767 | 8784266 | 10011855 |
| 1.25x | overlap | 14160480 | 14098034 | 9073516 | 9747395 | 10105258 |
| | no overlap | 12156673 | 12974662 | 6833634 | 7908482 | 9042862 |
| 1.5x | overlap | 15225117 | 15186672 | 9625059 | 10370551 | 10801828 |
| | no overlap | 11092036 | 11886024 | 6282091 | 7285326 | 8346292 |
| 1.75x | overlap | 16011058 | 15999422 | 10002613 | 10811681 | 11307364 |
| | no overlap | 10306095 | 11073274 | 5904537 | 6844196 | 7840756 |
| 2x | overlap | 16600114 | 16620928 | 10262561 | 11120285 | 11666168 |
| | no overlap | 9717039 | 10451768 | 5644589 | 6535592 | 7481952 |

Table 3: The number of markers considered for performance evaluation for each high-coverage sample and coverage level. The first row specifies the number of markers at which the filtered HQ data overlaps with the loci used in the imputation; these are the sites for which genotype concordance can be calculated. Subsequent rows show, for each coverage level that the high-coverage data was downsampled to, how many of the total sites had overlapping reads in the low-coverage data, and how many did not.

### References


[1] Iosif Lazaridis et al. "Ancient human genomes suggest three ancestral populations for present-day Europeans". In: *Nature* 513.7518 (Sept. 2014), pp. 409–413.

[2] Cristina Gamba et al. "Genome flux and stasis in a five millennium transect of European prehistory". In: *Nature Communications* 5 (Oct. 2014), p. 5257.

[3] Torsten Günther et al. "Population genomics of Mesolithic Scandinavia: Investigating early postglacial migration routes and high-latitude adaptation". In: *PLOS Biology* 16.1 (Jan. 2018), pp. 1–22.

[4] Lara M. Cassidy et al. "Neolithic and Bronze Age migration to Ireland and establishment of the insular Atlantic genome". In: *Proceedings of the National Academy of Sciences* 113.2 (2016), pp. 368–373.

[5] Pontus Skoglund et al. “Genomic diversity and admixture differs for Stone-Age Scandinavian foragers and farmers”. In: *Science* 344.6185 (2014), pp. 747–750.

[6] Rui Martiniano et al. “The population genomics of archaeological transition in west Iberia: Investigation of ancient substructure using imputation and haplotype-based methods”. In: *PLOS Genetics* 13.7 (July 2017), pp. 1–24.

[7] Iñigo Olalde et al. “A common genetic origin for early farmers from Mediterranean Cardial and Central European LBK cultures”. In: *Molecular Biology and Evolution* 32.12 (Sept. 2015), pp. 3132–3142.

[8] Andreas Keller et al. “New insights into the Tyrolean Iceman’s origin and phenotype as inferred by whole-genome sequencing”. In: *Nature Communications* 3.1 (2012), p. 698.

[9] Zuzana Hofmanová et al. “Early farmers from across Europe directly descended from Neolithic Aegeans”. In: *Proceedings of the National Academy of Sciences of the United States of America* 113.25 (June 2016), pp. 6886–6891.

[10] Torsten Günther et al. “Ancient genomes link early farmers from Atapuerca in Spain to modern-day Basques”. In: *Proceedings of the National Academy of Sciences* 112.38 (2015), pp. 11917–11922.

[11] Cristina Valdiosera et al. “Four millennia of Iberian biomolecular prehistory illustrate the impact of prehistoric migrations at the far end of Eurasia”. In: *Proceedings of the National Academy of Sciences* 115.13 (2018), pp. 3428–3433.

[12] Federico Sánchez-Quinto et al. “Megalithic tombs in western and northern Neolithic Europe were linked to a kindred society”. In: *Proceedings of the National Academy of Sciences* 116.19 (2019), pp. 9469–9474.

[13] Iain Mathieson et al. “Genome-wide patterns of selection in 230 ancient Eurasians”. In: *Nature* 528 (Nov. 2015), pp. 499–503.

[14] Eppie R. Jones et al. “Upper Palaeolithic genomes reveal deep roots of modern Eurasians”. In: *Nature Communications* 6.1 (2015), p. 8912.

[15] Gloria González-Fortes et al. “Paleogenomic evidence for multi-generational mixing between Neolithic farmers and Mesolithic hunter-gatherers in the lower Danube basin”. In: *Current Biology* 27.12 (June 2017), pp. 1801–1810.

[16] Alissa Mittnik et al. “The genetic prehistory of the Baltic Sea region”. In: *Nature Communications* 9.1 (2018), p. 442.
